# Internal state configures olfactory behavior and early sensory processing in *Drosophila* larvae

**DOI:** 10.1101/2020.03.02.973941

**Authors:** Katrin Vogt, David M. Zimmerman, Matthias Schlichting, Luis Hernandez-Nunez, Shanshan Qin, Karen Malacon, Michael Rosbash, Cengiz Pehlevan, Albert Cardona, Aravinthan D.T. Samuel

## Abstract

Animals exhibit different behavioral responses to the same sensory cue depending on their state at a given moment in time. How and where in the brain are sensory inputs combined with internal state information to select an appropriate behavior? Here we investigate how food deprivation affects olfactory behavior in *Drosophila* larvae. We find that certain odors reliably repel well-fed animals but attract food-deprived animals. We show that feeding state flexibly alters neural processing in the first olfactory center, the antennal lobe. Food deprivation differentially modulates two separate output pathways that are required for opposing behavioral responses. Uniglomerular projection neurons mediate odor attraction and show elevated odor-evoked activity in the food-deprived state. A multiglomerular projection neuron mediates odor aversion and receives odor-evoked inhibition in the food-deprived state. The switch between these two pathways is regulated by the lone serotonergic neuron in the antennal lobe, CSD. Our findings demonstrate how flexible behaviors can arise from state-dependent circuit dynamics in an early sensory processing center.

## Introduction

Hunger influences decisions about food-related sensory cues in many animal species. Whereas well-fed individuals can afford to be selective, individuals facing starvation must look for any available source of nutrition (Wu et al., 2005; Inagaki et al., 2014; Crossley et al., 2018). Since odors are commonly used to locate and identify food, olfactory responses can likewise vary with feeding state. In adult *Drosophila*, food deprivation has been shown to modulate the presynaptic excitability of certain olfactory receptor neurons (ORNs) (Root et al., 2011; Ko et al., 2015) and also the activity of dopaminergic input neurons in a higher brain region (Tsao et al., 2018). In contrast, the fixed architecture of the antennal lobe, the first olfactory processing center in the *Drosophila* brain, is thought to provide generic formatting of sensory inputs for use by various downstream circuits (Kazama and Wilson, 2008; Olsen and Wilson, 2008; Olsen et al., 2010; Wilson, 2013).

We asked whether the *Drosophila* larva responds differently to odors after food deprivation, and how its reduced nervous system might integrate internal state information to produce flexible innate behaviors. The larva has only 21 ORNs per hemisphere, each expressing a unique receptor type and innervating a distinct glomerulus in the larval antennal lobe (lAL). The complete wiring diagram of the lAL has been mapped by serial-section electron microscopy, providing a framework for understanding the algorithmic basis of larval olfaction (Berck et al., 2016). In addition to GABAergic broad local interneurons, well-studied in the adult (Olsen and Wilson, 2008), the lAL connectome also contains glutamatergic “picky” local interneurons (pLNs), whose function is as yet unknown. Another striking but poorly understood feature of the lAL is the existence of two types of projection neurons targeting different brain areas. Uniglomerular projection neurons (uPNs) relay signals from individual ORNs to the mushroom body calyx for associative learning and to the lateral horn for innate olfactory processing (Masuda-Nakagawa et al., 2009). Multiglomerular projection neurons (mPNs) receive input from different subsets of ORNs and innervate a wide variety of brain regions (Berck et al., 2016). Similar mPNs also exist in the adult fly, but they mainly project to the lateral horn (Tanaka et al., 2012; Bates et al., 2020). The lAL also contains a single, prominent serotonergic neuron (CSD) that is common to the larvae and adults of many insect species (Kent et al., 1987; Kloppenburg et al., 1999; Python and Stocker, 2002; Dacks et al., 2006; Roy et al., 2007). This surprising diversity of cell types hints at additional unknown computational functions.

We discovered that the larva’s behavioral response to an odor depends on its feeding state. For example, geranyl acetate (GA), which is innately aversive to fed larvae, becomes attractive to food-deprived larvae. This drastic change in behavioral response arises within the lAL circuitry. We observed no modulation of odor-evoked ORN responses, in contrast to what is seen in the adult. However, the two projection neuron output pathways show opposite state-dependent changes in odor-evoked activity and promote opposite innate behavioral responses. We found that food deprivation leads to an increase in odor-evoked CSD activity. Serotonergic signals from CSD directly excite the uPN pathway, which promotes odor attraction. Serotonin from CSD also recruits local glutamatergic inhibition and thereby indirectly suppresses the mPN pathway responsible for odor avoidance in the fed state. In summary, we reveal how statedependent neuromodulation reconfigures early odor processing by shifting the excitatory–inhibitory balance between two separate output pathways, thus allowing for execution of opposite behavioral responses to exactly the same sensory input.

## Results

### Feeding state determines the response to an odor

Food deprivation can alter food choice behaviors. A starving animal might approach food cues that a fed animal would avoid or ignore. To test this possibility in *Drosophila* larvae, we investigated olfactory behavior under different feeding states. Most monomolecular odorants are innately attractive to fed larvae (Fishilevich et al., 2005; Kreher et al., 2008). We screened a panel of 21 odorants and found two, geranyl acetate (GA) and menthol, that repel fed larvae but attract food-deprived larvae (Fig. 1A/B; S1A/B). Both GA and menthol occur naturally as antiherbivore compounds in green leaves (Bernasconi et al., 1998; Lin et al., 2017), which are not a preferred food source for fly larvae. We also tested the behavioral response to ethyl acetate (EA), a fruit odorant that is innately attractive to fed larvae (Fishilevich et al., 2005). While we did not observe a change in the sign of the behavioral response, EA becomes significantly more attractive after food-deprivation (Fig. S1C).

**Figure 1.**
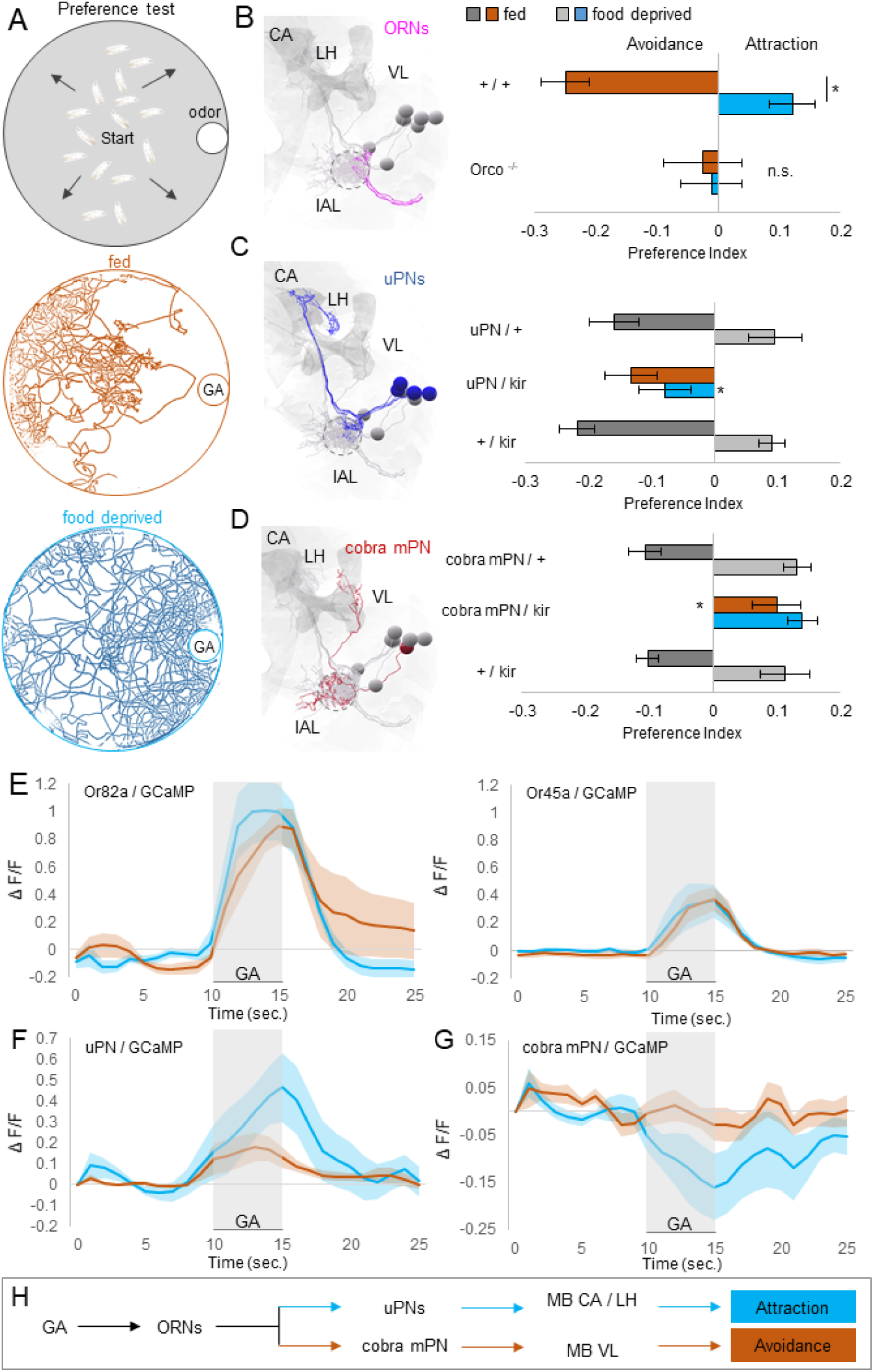
Feeding state determines the behavioral response to GA by modulating different lAL output pathways in opposite directions. **A** Behavioral setup to test olfactory preference. 15 larvae start in the middle of the arena and are free to move in the presence of an odor source. In the fed state, larvae avoid GA. Food deprived larvae are attracted to GA. Colored lines represent all larval tracks over 15 min. **B** EM reconstruction of GA ORNs (pink). Larvae show significant avoidance to GA in fed state (one sample *t*-test, *p <* 0.001) and significant attraction to GA in food deprived state (one sample *t*-test, *p <* 0.01). They show a significant switch from aversion to attraction upon food deprivation (two-sample *t*-test, *p <* 0.001). Mutant larvae (*Orco*^−*/*−^) that lack functional ORNs do not respond to the odor in any state (one sample *t*-test, *p >* 0.05) (*n* = 12–18). **C** EM reconstruction of GA uPNs (blue). Blocking neuronal output of uPNs (*GH146-GAL4*) by expressing *UAS-Kir2*.*1* impairs the food deprived (one-way ANOVA, post-hoc pairwise comparison, *p <* 0.05) but not fed (one-way ANOVA, *p >* 0.05) odor response. (*n* = 8–10). **D** EM reconstruction of cobra mPN (red). Blocking neuronal output of the cobra mPN by expressing *UAS-Kir2*.*1* under the control of *GMR32E03-GAL4* impairs avoidance in fed state (one-way ANOVA, post-hoc pairwise comparison, *p <* 0.001), but not attraction in food deprived state (one-way ANOVA, *p >* 0.05). (*n* = 6–10). **E** Calcium activity of the Or82a/Or45a neuron in response to GA (10^−6^) in fed and food deprived larvae. Odor was presented for 5 s. ORNs show the same activity levels in both states (*n* = 5–6).**F** Calcium activity in the uPNs in response to GA (10^−6^). Odor was presented for 5 s. uPNs show weak response in the fed state and an increased response after food deprivation (*n* = 7–9). **G** Calcium activity in the cobra mPN in response to GA (10^−6^). Odor was presented for 5 s. The cobra mPN shows decreased activity in the food deprived state (*n* = 8–9). **H** Two separate lAL output pathways receive opposing state dependent modulation and are required for opposing olfactory behaviors, respectively. Bar graphs represent pooled data from 5 min to 15 min during testing (mean ± SEM). Abbreviations: CA, mushroom body calyx; lAL, larval antennal lobe; LH, Lateral Horn; VL, mushroom body vertical lobe.

Mutant larvae (*Orco*^−*/*−^) lacking functional ORNs showed no response to GA, menthol, or EA when either fed or food-deprived (Fig. 1B; S1B/C). The state-dependent change in the behavioral response to these odorants requires the olfactory system. As every odorant activates different subsets of ORNs and other cell types in the lAL (Si et al., 2019), we focused our study on a single odorant, GA, to facilitate precise dissection of the mechanisms underlying the observed behavioral switch. Importantly, the transition from GA avoidance to attraction after 5-7 hours of food deprivation is highly reproducible across different odorant concentrations and larval stages (Fig. S2A-C).

### Feeding state modulates different lAL output pathways in opposite ways

The larval ORNs most sensitive to GA are those that express Or82a and Or45a (Si et al., 2019). What are the downstream targets of these sensory neurons? Each ORN connects to one uPN that innervates the mushroom body calyx and lateral horn (Fig. 1C). The *GH146-GAL4* line (Masuda-Nakagawa et al., 2009) drives expression in all uPNs activated by GA (Fig. S2E). From the wiring diagram of the lAL, we also identified an mPN called “cobra” that receives direct synaptic input from several ORNs including those activated by GA (Fig. 1D). Cobra mPN projects to the vertical lobe of the mushroom bodies (Berck et al., 2016). We also identified a driver line (*GMR32E02-GAL4*) specific for cobra mPN expression (Fig. S2F).

To probe the involvement of uPNs and cobra mPN in the innate behavioral response to GA, we inactivated each pathway by expression of Kir2.1, an inwardly rectifying potassium channel (Johns et al., 1999; Baines et al., 2001). Inactivation of uPNs made food-deprived larvae unable to switch their behavioral responses from avoidance to attraction (Fig. 1C). In contrast, inactivation of cobra mPN caused fed larvae to be attracted to GA without affecting attraction in food-deprived larvae (Fig. 1D).

To determine how feeding state affects odor-evoked activity of GA-sensing ORNs, uPNs, and the cobra mPN, we next recorded calcium dynamics by imaging GCaMP in intact, immobilized larvae (Si et al., 2019). First, we quantified odor-evoked activity in axon terminals of ORN-Or82a, ORN-Or45a or in all ORNs (Fig. 1E; S2D for statistics). We found no change in odor-evoked ORN activity after food-deprivation. In contrast, we found that food deprivation both increased odor-evoked uPN responses and inhibited cobra mPN (Fig. 1F/G; S2E/F for statistics).

These observations suggest that olfactory behavior might be computed within the antennal lobe circuit by changing how the same ORN input leads to different responses of the uPN and mPN output pathways (Fig. 1H). Higher uPN and lower mPN activity correlates with GA attraction. Lower uPN and higher mPN activity correlates with GA aversion.

### The mPN output pathway receives glutamatergic inhibition in food-deprived larvae

We sought synaptic mechanisms within the lAL that might upregulate the uPN pathway and downregulate the cobra mPN pathway. The glutamatergic picky local interneurons (pLNs) are well positioned to downregulate the mPN pathway because they preferentially synapse onto mPNs (Berck et al., 2016). Moreover, GluCl*α*, a glutamate-gated chloride channel that makes glutamate inhibitory, is widely expressed in the adult antennal lobe (Liu and Wilson, 2013).

When we reduced GluCl*α* expression specifically in cobra mPN, both fed and food-deprived animals exhibited GA avoidance (Fig. 2A). Similar to what we observed in previous imaging experiments with wild-type animals, food deprivation decreased odor-evoked mPN responses. However, after GluCl*α* knockdown, we found no change in GA-evoked calcium dynamics between fed and food-deprived animals (Fig. 2B/C; S3A for statistics). Removing glutamatergic inhibition of cobra mPN appears to prevent inactivation of the odor avoidance pathway after food deprivation and odor response does not shift to attraction.

**Figure 2.**
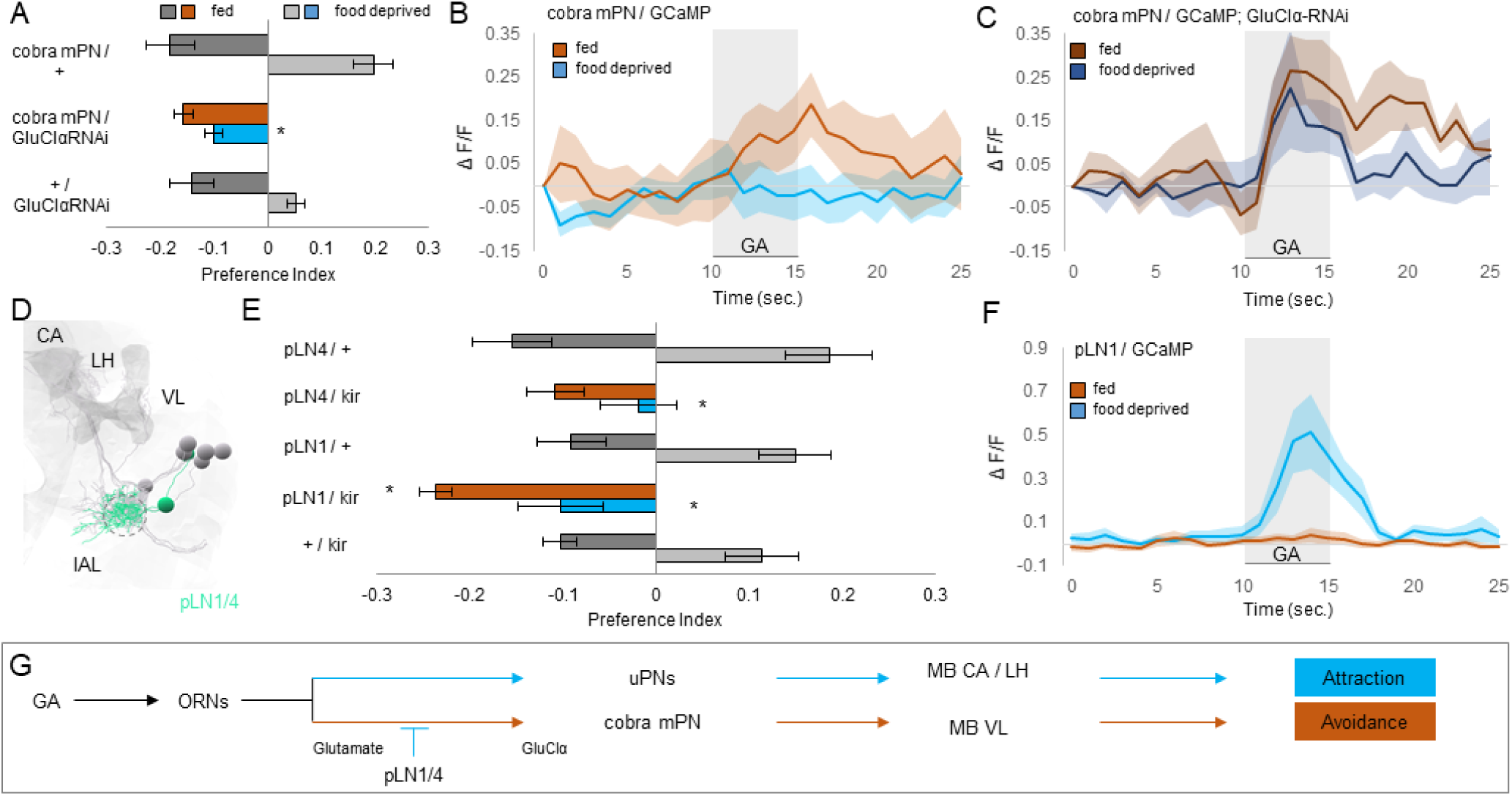
Upon food deprivation, the mPN pathway receives glutamatergic inhibition from pLNs. **A** Knockdown of the GluCl*α*-receptor using RNAi in cobra mPN has no effect on odor avoidance in the fed state (Kruskal-Wallis test, *p >* 0.05). Odor attraction in the food-deprived state is impaired (Kruskal-Wallis test, post-hoc pairwise comparison, *p <* 0.001) (*n* = 4–8). **B, C** Calcium activity in the cobra mPN in response to GA (10^−5^). Odor was presented for 5 s. Light colors: The cobra mPN shows decreased activity in the food-deprived state. Dark colors: When expressing GluCl*α* RNAi, we cannot detect any state dependent inhibition (*n* = 8). **D, E** EM reconstruction of local lAL interneurons, pLN1/4 (bright green). Blocking output of pLN1 slightly increases avoidance in the fed state (Kruskal-Wallis test, post-hoc comparison, *p <* 0.05 (pLN1), *p >* 0.5 (pLN4). Blocking pLN1 and 4 leads to loss of attraction in the food-deprived state (Kruskal-Wallis test, post-hoc comparison, *p <* 0.05). (*n* = 8–10). **F** Calcium activity in pLN1 in response to GA (10^−6^). Odor was presented for 5 s. The pLN1 shows increased activity in the food-deprived state (*n* = 7). **G** pLNs are glutamatergic and provide inhibition onto cobra mPN by GluCl*α*-receptor binding in the lAL. Bar graphs represent pooled data from 5 min to 15 min during testing (mean ± SEM). Abbreviations: CA, mushroom body calyx; lAL, larval antennal lobe; LH, Lateral Horn; VL, mushroom body vertical lobe.

There are four glutamatergic picky local interneurons (pLNs) that receive input from GA-sensitive ORNs and send output to the cobra mPN: pLN0, pLN1, pLN2, and pLN4 (Berck et al., 2016). The wiring diagram only contains one other pLN (pLN3), which shares synapses with neither the GA-sensitive ORNs nor cobra mPN. We identified three cell-specific split GAL4 lines for pLN1, pLN3, and pLN4 (Fig. 2D; S3B-G). Targeted inactivation of pLN1, pLN3, or pLN4 had no effect on GA avoidance in fed larvae, but inactivating either pLN1 or pLN4 eliminated GA attraction in food-deprived larvae (Fig. 2E; S3H). With imaging, we found that GA-evoked calcium responses in pLN1 were elevated in the food-deprived state and undetectable in the fed state (Fig. 2F; S3I for statistics). We conclude that food deprivation may downregulate the cobra mPN pathway by increasing glutamatergic inhibition from pLN1 and pLN4 (Fig. 2G).

### uPNs receive serotonergic excitation through the 5-HT7 serotonin receptor

How might the activity of the uPN pathway be upregulated in food-deprived animals? Serotonin can be a prominent neuromodulator, and the excitatory 5-HT7 receptor is expressed in the uPN pathway (Huser et al., 2017; Saudou et al., 1992) (Fig. 3A; S4A). We found that inactivating 5-HT7 expressing neurons did not impair GA avoidance in fed animals, consistent with previous observations (Huser et al., 2017) (Fig. 3B). However, inactivating 5-HT7 expressing neurons caused food-deprived larvae to avoid GA, similar to the phenotype obtained by inactivating all uPNs. We also specifically removed 5-HT7 from the uPN pathway using a CRISPR/Cas9-based cell type-specific gene knockout system (Schlichting et al., 2019; Delventhal et al., 2019). Without 5-HT7 in the uPNs, food-deprived larvae also avoided GA (Fig. 3C). With imaging, we found that odor-evoked calcium activity in the uPNs lacking 5-HT7 was weak in both food-deprived and fed animals and did not increase after food deprivation (Fig. 3D/E; S4B for statistics). We conclude that serotonin is required to elevate odor-evoked uPN activity and to switch the behavioral response to attraction in food-deprived animals.

**Figure 3.**
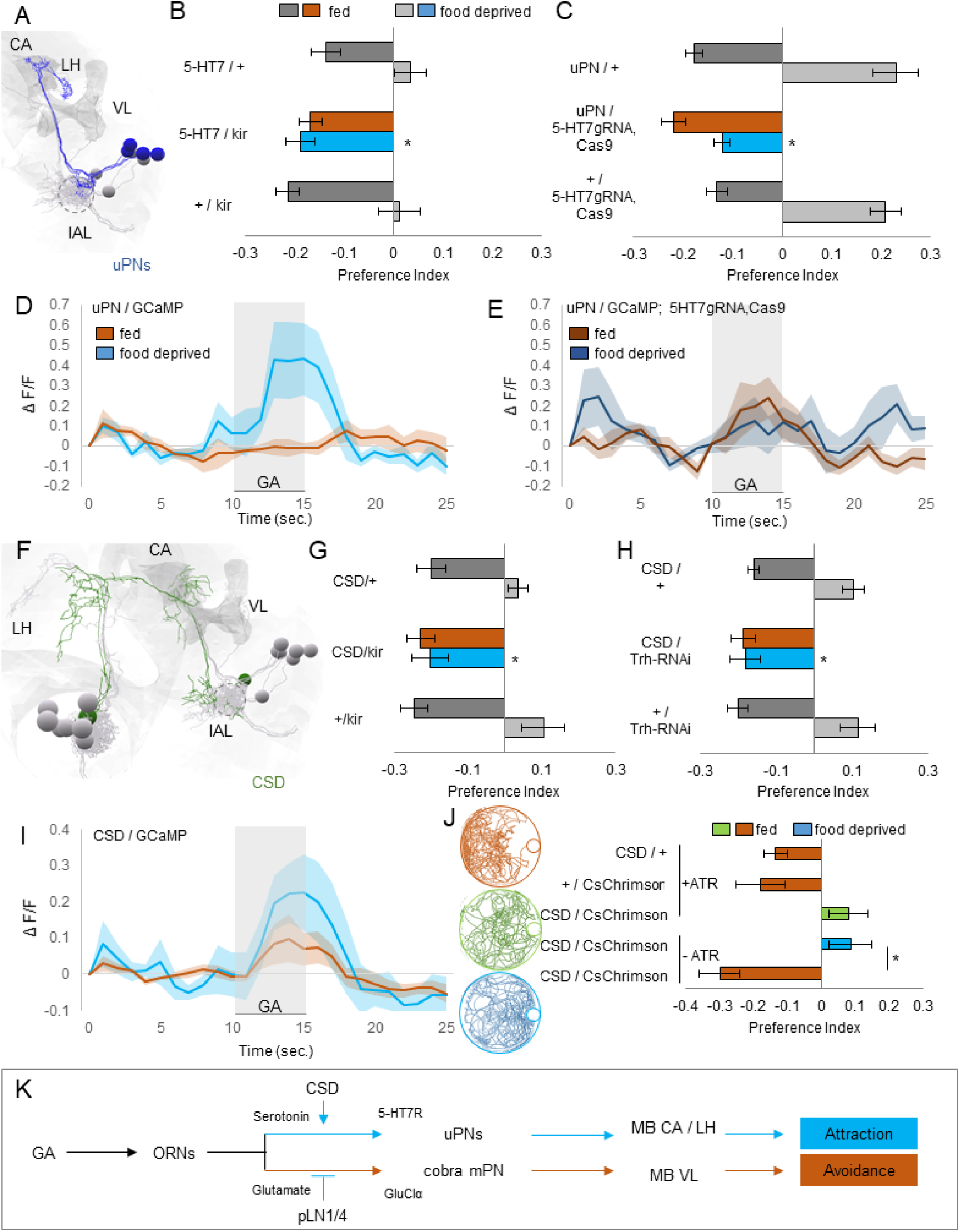
Upon food deprivation, CSD elevates uPN activity via the excitatory 5-HT7 receptor. **A, B** EM reconstruction of uPNs (blue). Blocking neuronal output of 5-HT7 receptor-expressing neurons (*5-HT7-GAL4*, including uPNs) with *UAS-Kir2*.*1* does not affect odor avoidance in fed larvae (one-way ANOVA, *p >* 0.05); however, odor attraction in food-deprived larvae is impaired (one-way ANOVA, post-hoc pairwise comparison, *p <* 0.01). (*n* = 8). **C** Knockdown of the 5-HT7 receptor with *UAS-Cas9* in the uPNs (*GH146-GAL4*) does not affect odor avoidance in fed larvae (one-way ANOVA, post-hoc pairwise comparison, *p >* 0.05), however odor attraction in food-deprived larvae is impaired (one-way ANOVA, post-hoc pairwise comparison, *p <* 0.001) (*n* = 8–14). **D**,**E** Calcium activity in the uPNs in response to GA (10^−6^). Odor was presented for 5 sec. Light colors: The uPNs show increased activity in the food-deprived state. Dark colors: When knocking down the 5-HT7 receptor with *UAS-Cas9* in the uPNs, we cannot detect any state dependent increase in response (*n* = 6–7). **F**,**G** EM reconstruction of the CSD neuron (dark green). Blocking neuronal output of the CSD neuron (*R60F02-GAL4*) via expression of *UAS-Kir2*.*1*, we find no effect on behavior in the fed state (one-way ANOVA, *p >* 0.05). However, food-deprived larvae do not switch their behavior towards attraction (Kruskal-Wallis test, post-hoc pairwise comparison, *p <* 0.05). (*n* = 6–8). **H** Preventing serotonin synthesis in the CSD neuron via expression of *UAS-Trh-RNAi* does not affect odor avoidance in fed larvae (Kruskal-Wallis test, *p >* 0.05). however odor attraction in food-deprived larvae is impaired (Kruskal-Wallis test, post hoc pairwise comparison, *p <* 0.01). (*n* = 8). **I** Calcium activity in the CSD neuron (*R60F02-GAL4*) in response to GA (10^−8^). Odor was presented for 5 sec. The CSD neuron responds stronger to GA in the food-deprived state (*n* = 8–10). **J** Optogenetic activation of the CSD neuron impairs odor avoidance in fed larva (one-way ANOVA, post-hoc pairwise comparison, *p <* 0.05) (*n* = 12–20). **K** Higher activity and serotonin release by the CSD neuron lead to higher activation of the uPNs via the excitatory 5-HT7 receptor. This induces odor attraction in the food-deprived state. Bar graphs represent pooled data from 5 min to 15 min during testing (mean ± SEM). Abbreviations: CA, mushroom body calyx; lAL, larval antennal lobe; LH, Lateral Horn; VL, mushroom body vertical lobe.

### Serotonin released from CSD induces a state-dependent switch in odor response

A single serotonergic neuron in each hemisphere called CSD widely innervates higher brain areas as well as the antennal lobes in both the larva and adult fly (Dacks et al., 2006; Roy et al., 2007; Zhang and Gaudry, 2016; Huser et al., 2012) (Fig. 3F; S4C). We found that cell-specific inactivation of CSD using *R60F02-GAL4* (but not inactivation of other serotonergic neurons, Fig. S4D) causes GA avoidance in both fed and food-deprived larvae (Fig. 3G). This phenotype was also obtained by cell-specific knockdown of serotonin synthesis in CSD and exhibited by serotonin-synthesis mutants (Fig. 3H; S4E).

CSD receives a few synapses from ORNs, but also substantial indirect olfactory input from neurons in the lateral horn that integrate signals from uPNs (Berck et al., 2016). We measured GA-evoked calcium dynamics in CSD, and found an increase in food-deprived animals compared to fed animals (Fig. 3I; S4F for statistics). To determine whether elevated CSD activity is linked to a change in behavior, we used optogenetics. We expressed the red-shifted optogenetic effector CsChrimson in CSD and tested behavior. Fed larvae that normally show GA avoidance will tend to exhibit attraction when CSD is artificially activated by continuous illumination with red light (Fig. 3J; S5A). We note that CSD activation does not affect basal locomotion patterns (e.g., lengths of crawling movements) in the same way as food deprivation (Fig. S5B). CSD activation changes olfactory responses, but not the ability to crawl towards an olfactory cue.

Inhibition of the serotonin transporter SerT, which localizes in presynaptic membranes and recycles released neurotransmitter, is a method for increasing serotonergic neurotransmission (Giang et al., 2011; Xu et al., 2016). RNAi knockdown of SerT in CSD reduced GA avoidance in fed larvae (Fig. S4G). In contrast, overexpression of SerT (which presumably lowers serotonergic transmission) causes GA avoidance in both fed and food-deprived larvae (Fig. S4G). These phenotypes are consistent with optogenetic activation of CSD and constitutive inactivation of CSD, respectively. These phenotypes also argue against the possibility that state-dependent changes in receptor expression occur in the projection neurons themselves.

Notably, the wiring diagram reveals no direct synapses from CSD to the uPNs (Fig. 4A/B). One possibility is that 5-HT7 receptors in the uPNs might be activated by extrasynaptic release or synaptic spillover after food deprivation (Trueta and De-Miguel, 2012; De-Miguel and Trueta, 2005) (Fig. 3K).

**Figure 4.**
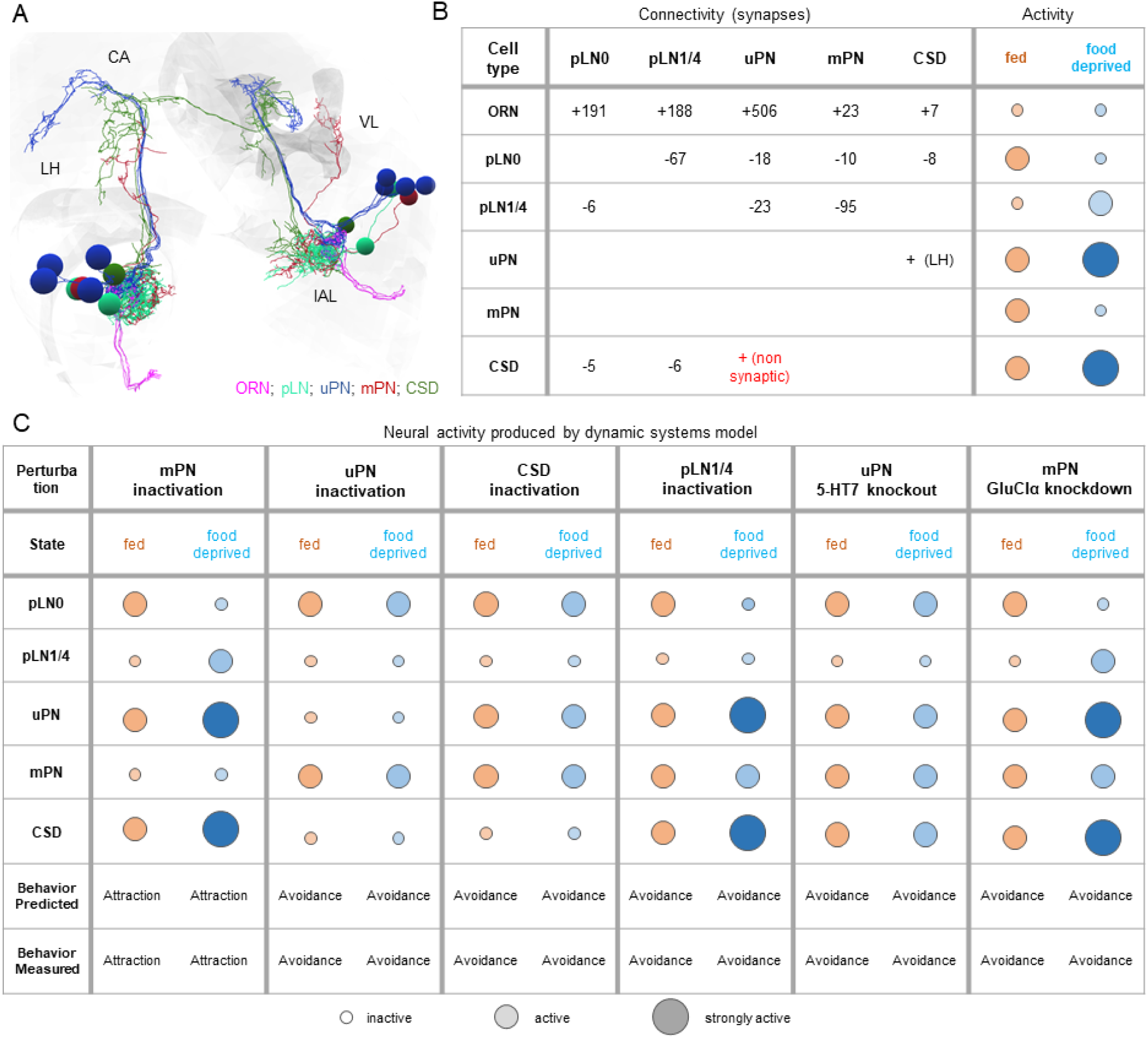
A simple dynamical model recapitulates state-dependent changes in odor valence. **A** EM reconstructions of all studied neurons in the larval brain. The right mushroom body is shaded gray. Abbreviations: CA, mushroom body calyx; lAL, larval antennal lobe; LH, Lateral Horn; VL, mushroom body vertical lobe. **B** EM connectivity of all studied neurons. Number of synapses between cell types. Circles indicate neural activity of all studied lAL neurons based on functional imaging recordings and behavioral experiments (fed [orange] vs. food-deprived [blue]). In the fed state, the cobra mPN pathway, required for odor avoidance, is active, because cobra mPN does not receive inhibition from pLN1/4s. Upon food deprivation, serotonergic modulation mediated by CSD increases and uPNs receive extrasynaptic modulation via the excitatory 5-HT7 receptor. The mPN receives glutamatergic inhibition from pLN1/4s. **C** Neural activity of all studied lAL neurons in fed (orange) and food-deprived state (blue) upon cell manipulations within the dynamical systems model. Bottom rows: Behavioral output is calculated based on activity of the uPNs and the mPN. The model predicts the same behavioral output as we found in behavioral experiments.

### Food-deprivation modulates inhibitory interactions in the pLN circuit

Our results suggest that a pLN circuit regulates mPN activity by providing glutamatergic inhibition in the food-deprived state. But what inactivates pLN1/4 in the fed state? The wiring diagram suggests that pLNs inhibit one another: pLN1/4 receive strong glutamatergic inhibition from pLN0 (Berck et al., 2016) (Fig. 4B). We were not able to study pLN0 due to lack of a cell-specific driver. However, we found that reducing GluCl*α* expression in pLN1 and pLN4 eliminates GA avoidance in the fed state (Fig. S6A/B). Thus, pLN1/4 receive glutamatergic inhibition, presumably from pLN0, only when larvae are fed.

Additionally, the local interneurons of the lAL express an inhibitory serotonergic receptor, 5-HT1A (Huser et al., 2017) (Fig. S6C). CRISPR-mediated knockout of the 5-HT1A receptor from pLN1/4 similarly abolished GA avoidance in the fed state (Fig. S6D). According to the wiring diagram, pLN1/4 receive direct synaptic inputs from CSD that seem to provide enough serotonin in the fed state to achieve this inhibition (Fig. 4B). Thus, in the fed state, pLN1/4 receive joint inhibition from glutamatergic and serotonergic signals, thereby lowering their ability to inhibit the mPN pathway for odor avoidance (Fig. S6E).

In the food-deprived state, knockdown of GluCl*α* in pLN1/4 does not affect odor attraction. This suggests that they receive less glutamatergic inhibition (presumably from pLN0) and are thus able to downregulate the mPN pathway (Fig. S6A/B). Both glutamatergic and serotonergic inputs seem to be required for inhibition of pLN1/4 and the resulting disinhibition of the mPN pathway, as increased serotonergic inhibition from CSD in the food-deprived state appears to be insufficient to inactivate pLN1/4 (Fig. S6D). In contrast, inhibition of pLN0 is likely inrdependent of glutamate, as it lies exclusively upstream of the rest of the pLN network. Since all five pLNs originate from the same neuroblast lineage (Berck et al., 2016; Das et al., 2013), pLN0 is also likely to express the inhibitory 5-HT1A receptor and therefore be subject to inhibition from CSD under food deprivation.

### A computational model of the state-dependent switch in olfactory response

To integrate our findings about synaptic properties and state-dependent neuromodulatory dynamics with the known circuit connectivity, we built a computational model of the full lAL network (Fig. 4A/B). In particular, having uncovered the sign of the relevant synaptic and non-synaptic interactions, we set out to test whether the resulting dynamics would give rise to decision-making: namely the appropriate changes in activity in the uPN and mPN output pathways. In our model, the weight of every synaptic connection is calculated from the number of synapses between cell types in the connectome (Berck et al., 2016) (Fig. 4B). The only plausible cellular input to pLN1/4 that provides glutamatergic inhibition is pLN0, which, like pLN1/4, is modeled as receiving direct serotonergic inhibition from CSD.

Our model, based on the established connectivity and empirically determined synaptic properties, recapitulates the observed changes in neuronal activity. Activation of CSD in the food-deprived state upregulates the uPN pathway directly through serotonergic excitation as well as inhibits the mPN pathway indirectly through recruitment of glutamatergic pLNs (Fig. S7). In contrast, weak CSD activity in the fed state shifts activity towards the mPN pathway.

Our model also allows us to test the consequences of all the molecular and cellular perturbations that we performed to uncover circuit properties. We simulated the effects of removing individual neurons from the circuit (akin to chronic inactivation by Kir2.1) as well as individual synapses (akin to receptor knockdown or knockout), and found that, in every case, we could predict whether the circuit output would be shifted towards avoidance or attraction in either fed or food-deprived animals (Fig. 4C). Taken together, these data suggest that our minimal circuit model is sufficiently comprehensive to reproduce the key state-dependent changes.

## Discussion

The insect antennal lobe, like the mammalian olfactory bulb, is typically regarded as a preprocessing stage whose basic function is to format inputs in a generic way for use by various downstream circuits (Wilson, 2013). In contrast, we have demonstrated a crucial role for the lAL in actively implementing a state-dependent behavioral switch between avoidance and attraction to certain odors. In particular, we show that food deprivation reverses the larva’s innate aversion to GA. We also reproduce our key observations about GA using menthol, the only other odorant that we found to elicit consistent avoidance behavior in well-fed larvae (Fig. S8A-G). Both odorants are monoterpenoids, a class of volatile organic compounds that plants secrete to defend against herbivory (Rice and Coats, 1994). Nevertheless, it may be adaptive for larvae to suppress their default aversion to these potential toxins when faced with starvation.

Remarkably, we find that this behavioral choice is implemented by a state-dependent shift in the relative activity of two differentially projecting lAL output pathways. In the adult fly, both the uPNs and most mPNs project to the lateral horn, where their activity is integrated to generate innate behaviors (Strutz et al., 2014; Wang et al., 2014). In the larva, uPNs project to the mushroom body calyx and lateral horn and promote attractive behavior, while the previously uncharacterized cobra mPN projects to the mushroom body vertical lobe and promotes aversive behavior. By regulating the state-dependent switch between these pathways, the lAL contributes to the assignment of innate odor valence. Understanding how the uPNs and mPN organize locomotory behavior toward or away from odors will require mapping downstream pathways to the motor circuit (Tastekin et al., 2015, 2018). However, we find that the key neuromodulatory switch between these output pathways is implemented within the lAL, which must therefore be regarded as a *bona fide* decision-making circuit.

Serotonergic neuromodulation has been implicated in numerous state-dependent behaviors—such as aggression, circadian entrainment, olfactory learning, and feeding (Johnson et al., 2009; Nichols, 2007; Neckameyer et al., 2007; Schoofs et al., 2018; Tsao et al., 2018)—in both vertebrates and invertebrates (Dayan and Huys, 2009; Bacqué-Cazenave et al., 2020). We find that CSD, a prominent serotonergic neuron whose processes span the *Drosophila* olfactory system, controls the switch between attraction and aversion to GA through state-dependent modulation of the uPN and mPN output pathways. CSD exhibits greater odor-evoked activity in the food-deprived state than in the fed state. When an animal is food-deprived, high levels of serotonin released from CSD both activate the uPN pathway (via the excitatory 5-HT7 receptor) and increase glutamatergic inhibition onto the mPN pathway (via modulation of inhibitory glutamatergic local interneurons). As there are no direct synapses from CSD onto the uPNs, serotonergic transmission likely involves either synaptic overspill or non-synaptic neurotransmitter release (De-Miguel and Trueta, 2005). Serotonin from CSD has also been shown to excite uPNs in the adult fly (Dacks et al., 2009). However, the adult CSD neuron exhibits more elaborate patterns of innervation (Coates et al., 2017) as well as compartmentalized odor-evoked responses (Zhang et al., 2019). Moreover, each cell type in the adult antennal lobe expresses multiple types of serotonin receptors, suggesting the possibility for crosstalk (Sizemore and Dacks, 2016). By exploiting the numerical simplicity of the lAL, we have succeeded in unravelling the neuromodulatory mechanism that generates flexible behavior from a fixed wiring diagram.

The neural circuit for early olfactory processing in mammals, the olfactory bulb, is strikingly similar in its molecular and circuit architecture to the *Drosophila* antennal lobe (Gaudry, 2018). Here, we have shown that CSD activity encodes information about feeding state which the lAL uses to select an appropriate behavioral response. In the mouse, serotonergic projection neurons from the raphe nucleus innervate the olfactory bulb (McLean and Shipley, 1987) and also modulate distinct local interneurons and output pathways (Kapoor et al., 2016; Brunert et al., 2016). The circuit logic and behavioral role of serotonergic signaling in mammalian olfaction is not well understood. However, homologous 5-HT receptors are known to regulate appetite and seeking/craving behaviors, suggesting a conserved function for serotonergic regulation of behaviors that depend on feeding state (Ebenezer et al., 2007; Hauser et al., 2015). By systematically analyzing the circuit-level effects of state-dependent neuromodulation in the lAL, our study suggests a potentially general mechanism by which internal state can modify early sensory processing to determine behavior.

## ACKNOWLEDGEMENTS

We would like to thank members of the Samuel and de Bivort labs for helpful discussions. We also thank Yvette Fisher, Helen Yang, and other members of the Wilson lab; Margaret Herre and other members of the Vosshall lab; as well as Venkatesh Murthy, Armin Bahl, Dana Galili, Benjamin Kottler, and Kenny Blum for their comments on the manuscript. We acknowledge the Bloomington *Drosophila* Stock Center (NIH P40OD018537), Michael Pankratz and James Truman for fly lines. Brian Smith kindly provided odorants. Microfabrication was performed iCenter for Nanoscale Systems and the NSF’s National Nanotechnology Infrastructure Network (NNIN) K.V. was supported by a DFG research fellowship (project no. 345729665). C.P. acknowledges support by the NIH. A.D.T.S. is supported by grants from the NSF and NIH.

## AUTHOR CONTRIBUTIONS

K.V. performed behavioral experiments, functional imaging, anatomical studies and optogenetics. D.M.Z. performed functional imaging. M.S. and M.R. provided genetic reagents. K.M. performed behavioral experiments. L.H.N. analyzed data from optogenetic experiments. S.Q. and C.P. designed the computational model. A.C. provided access to connectomic data. K.V. conceived the project. K.V. and A.D.T.S. designed the study, interpreted the results, and wrote the manuscript with input from all authors.

## Methods

### Experimental model

Flies were reared at 22 °C under a 12:12 light-dark cycle and 60% humidity in vials containing standard cornmeal agar-based medium. For larval experiments, adult flies were transferred to larvae collection cages (Genesee Scientific) containing grape juice agar plates and 180 mg of fresh yeast paste per cage. Flies were allowed to lay eggs on the agar plate for 1–2 days before the plate was removed for collection of larvae in the different developmental stages. Behavioral experiments were performed with L2 larvae (3–4 days after egg laying), unless otherwise stated (Fig. S2A). Calcium imaging experiments and anatomical studies were performed in L1 larvae (2 days after egg laying). Most transgenic stocks were obtained from Bloomington *Drosophila* Stock Center (BDSC, see Table 1).

**Table 1.** Fly lines used in this study.

| Line | Source |
| --- | --- |
| *Orco*<sup>1</sup> | BDSC #23129 |
| *GH146-GAL4* | BDSC #30026 |
| *GMR32E03-GAL4* | BDSC #49716 |
| *UAS-Kir2.1* | BDSC #6596 |
| *Orco-GAL4* | BDSC #23292 |
| *Or82a-GAL4* | BDSC #23125 |
| *Or45a-GAL4* | BDSC #9975 |
| *UAS-GCaMP6m; Orco::RFP* | This study (*UAS-GCaMP6m*, BDSC #42748; *Orco::RFP*, BDSC #63045) |
| *UAS-GluClα-RNAi (TRiP HMC03585)* | BDSC #53356 |
| *pLN1-GAL4 (split)* | This study (*GMR21D06-AD*, BDSC #70117; *GMR50A06-DBD*, BDSC #68988) |
| *pLN3-GAL4 (split; SS004499)* | J. Truman (*GMR42E06-AD*, BDSC #71054; *GMR12C03-DBD*, BDSC #70429) |
| *pLN4-GAL4 (split; SS001730)* | J. Truman (*GMR21D06-AD*, BDSC #70117; *GMR12C03-DBD*, BDSC #70429) |
| *UAS-mCD8::GFP; Orco::RFP* | BDSC #63045 |
| *5-HT7-GAL4* | M. Pankratz (Becnel et al., 2011) |
| *UAS-gRNA-5-HT7; UAS-Cas9* | This study (Rosbash lab) |
| *GMR60F02-GAL4* | BDSC #48228 |
| *UAS-Trh-RNAi (TRiP JF01863)* | BDSC #25842 |
| *UAS-CsChrimson::mVenus* | BDSC #55136 |
| *Trh-GAL4* | BDSC #38389 |
| *DDC-GAL4* | BDSC #7009 |
| *Trh<sup>c01440</sup>* | BDSC #10531 |
| *UAS-SerT* | BDSC #24464 |
| *UAS-SerT-RNAi (TRiP HMJ30062)* | BDSC #62985 |
| *5-HT1A-GAL4* | M. Pankratz (Luo et al., 2012) |
| *UAS-gRNA-5-HT1A; UAS-Cas9* | This study (Rosbash lab) |
| *y<sup>1</sup>w<sup>67c23</sup>; P{CaryP}attP1* | BDSC #8621 |
| *w<sup>1118</sup>; sna<sup>Sco</sup> It<sup>1</sup>/CyO; MKRS/TM6B* | BDSC #3703 |

### Cell-specific CRISPR/Cas9-mediated knockout

We generated *UAS-gRNA* lines targeting the 5-HT7 and 5-HT1A receptors, as described previously (Port and Bullock, 2016). In short, we digested the pCFD6 vector (a gift from Simon Bullock, Addgene #73915) with BbsI (New England Biolabs, NEB) and used a Gibson Assembly (NEB) to incorporate PCR products. We generated two PCR fragments each harboring three guide sequences with homology to the CDS of either *5-HT7* or *5-HT1A* (see Table 2). The resulting colonies were sequenced and the correct constructs were inserted into the *attP1* landing site on chromosome II (BDSC #8621) by ΦC31-mediated recombination (Rainbow Transgenic Flies, Camarillo, CA, USA). Transgenic flies were back-crossed to *w*^1118^ and balanced using BDSC #3703 (Schlichting et al., 2019; Delventhal et al., 2019).

**Table 2.**
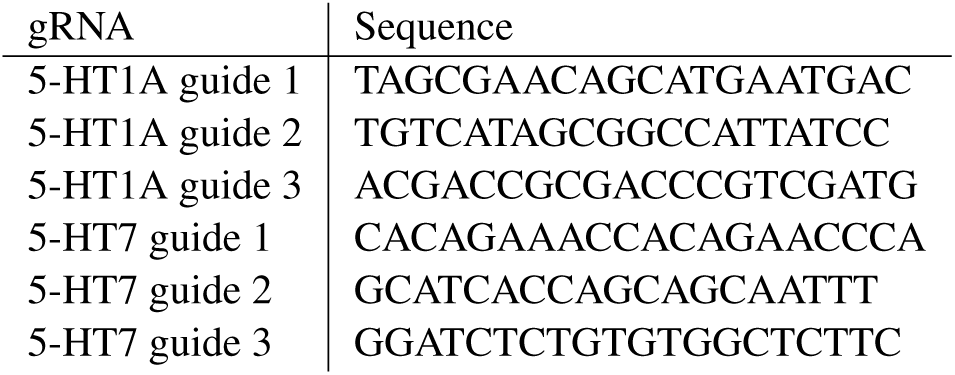
Guide sequences.

### Behavioral assays

Pure odorants were diluted in deionized (DI) water and stored for no more than one week. Our initial screen of behavioral responses in fed larvae involved an odorant panel of natural constituents of ripe fruit from Brian Smith (Keesey et al., 2015, based on) (Fig. S1A). Each odorant and odor dilution was stored in the dark and in a separate glass bottle to avoid contamination. For behavior experiments, the following odorants were obtained from Sigma-Aldrich and used at the indicated dilutions (v/v): geranyl acetate, 10^−4^ (except see Fig. S2B; CAS #105-87-3); menthol, 10^−3^ (CAS #89-78-1); ethyl acetate, 10^−6^ (CAS #141-78-6). The larval olfactory choice assay is illustrated in Fig. 1A. For the measurement of olfactory responses in the fed state, 3– 4-day-old larvae were picked from grape juice plates and briefly washed in several DI water droplets right before the experiment. For odor preference test in the food-deprived state, larvae were picked from grape juice plates 5–7 h before test (except as otherwise stated, Fig. S2C), washed in DI water droplets and placed in a small Petri dish (VWR, #60872-306, 60 mm *×* 15 mm) with a 2% agarose plate on the bottom covered by a thin layer of tap water. Before odor preference test, food-deprived larvae were placed in DI water droplets similar to larvae freshly picked from food. At the start of the test, 15 larvae were placed on the surface of a 2% agarose pad in the center of a 10 cm Petri dish equipped with a 12 mm plastic cup at one edge (VWR, #25384-318). Before each experiment, the cup was loaded with 200 µL of odor solution. Larvae were free to explore the arena with closed lid for 15 min. Over time, we counted the number of larvae in each quadrant of the arena, and computed a preference index:

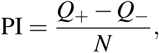

where *Q*_+_ represents the number of larvae in the quadrant containing the odor cup, *Q*_−_ represents the number of larvae in the opposite quadrant, and *N* represents the total number of larvae in the arena.

Each odor preference test used freshly picked and previously untested larvae. Different larvae were tested under fed and food-deprived conditions. Experiments were performed at room temperature under uniform illumination with a point light source (desk lamp) on a clean bench, except for the optogenetic experiments shown in Fig. 3J (see below). In all experiments, we alternated the position of the odor cup in the Petri dish between the left or right sides to avoid systematic bias.

### Optogenetics

For optogenetic experiments with CsChrimson, experimental larvae additionally received food supplemented with 0.1 mM all-trans-retinal (ATR; Sigma-Aldrich, CAS #116-31-4). Larval collection cages and grape juices plates were wrapped in foil and kept in dark during egg laying and larval development (Hernandez-Nunez et al., 2015). Experiments were performed in a lightproof box. Olfactory response behavior was recorded at 4 Hz using a CCD Mightex camera equipped with a long-pass infrared filter (cutoff 740 nm). Larvae received 500 Hz pulsed stimulation with spatially uniform red light (625 nm, pulse width 20 µs, 0.62 W*/*m^2^ ± 0.5 %) to activate the optogenetic effector throughout the entire 15 min test period. Larval trajectories were analyzed with custom tracking software (Ger-show et al., 2012). For each frame, the number of larvae in the two halves of the Petri dish was analyzed automatically. The preference index was calculated as

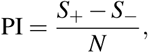

where *S*_+_ represents the number of larvae on the half side containing the odor cup, *S*_−_ represents the number of larvae on the opposite side, and *N* represents the total number of larvae in the arena. The PI over each min of the test was pooled for comparable representation of data in Fig. S5A.

### Functional imaging with microfluidic device

We used a previously described method for microfluidics and imaging (Si et al., 2019). An 8-channel microfluidic chip was used in this study to deliver odor solutions to an intact larva. The same odors were used as in the behavior setup, however at lower concentrations (GA: 10^−8^, 10^−6^, 10^−5^; menthol: 10^−4^). Stimuli consisted of 5 s odor pulses interleaved with 15 s water washout periods. L1 larvae were loaded into the microfluidic device using a 1 mL syringe filled with Triton X-100 (0.1% [v/v]) solution. The larva was pushed to the end of the loading channel with its dorsal side closest to the objective. GCaMP signal was recorded using an inverted Nikon Ti-E spinning disk confocal microscope with a 60X water immersion objective (NA 1.2). A CCD microscope camera (Andor iXon EMCCD) captured frames at 30 Hz. The CSD neuron and projection neurons were recorded by scanning the entire volume (step size 1.5 µm) of the brain, ranging from the antennal lobe to the processes in the higher brain. Orco::RFP was used to locate the lAL. Recordings from at least 5–9 larvae were collected for each genotype and condition. All samples were used for analysis unless dendritic varicosities developed in the ORNs over the course of the recording, a sign of unhealthy neurons likely due to mechanical stress.

### Anatomical studies

GFP expression patterns of GAL4 and split-GAL4 lines were imaged in intact larvae using an inverted Nikon Ti-E spinning disk confocal microscope with a 60X water-immersion objective (NA 1.2) or 20X objective. To immobilize larvae, they were squeezed between two glass slides using a micromanipulator. Orco::RFP was used to locate the lAL.

### Quantification and statistical analysis

Imaging data were analyzed with custom scripts written in MATLAB, available from GitHub. Data from behavioral experiments were analyzed with LabVIEW, MATLAB and Microsoft Excel. Data were tested for normality (Shapiro-Wilk test) and analyzed by parametric or non-parametric statistics as appropriate: the Kruskal-Wallis test or one-way analysis of variance (ANOVA). For post-hoc pairwise comparisons, the two-tailed one- or two-sample *t*-test, Welch’s *t*-test, or Mann-Whitney *U* - test were performed as appropriate. The significance level of statistical tests was set to *α* = 0.05. Only the most conservative statistical result of multiple pairwise comparisons is indicated. No statistical methods were used to determine sample sizes in advance, but sample sizes are similar to those reported in other studies in the field. Sample sizes (“*n*”), *p* values, and other relevant test statistics are shown in the appropriate figure legends. Bar graphs represent pooled data from 5 min to 15 min during testing (mean ± SEM). Box plots show single data points, median (50th percentile), and quartiles (25th/75th percentile) of data. Whiskers show maximum and minimum of data.

### Computational circuit model

To better understand how the neural circuitry in the lAL dictates the state-dependent shift of odor valence, we built a dynamical model based on connectomic data and neural activity of larval neurons. We assume that the neural activities for pLNs and cobra mPN are binary (0 and 1), while uPN and CSD neurons have three states, 0, 1, and 2. We denote the state of a neuron at time point *t* as *S*_*i*_(*t*) ∈ {0, 1, 2}, *i* = 1, …, 5. For simplicity, we have pooled pLN1 and pLN4 (pLN1/4). The synaptic weights and other interactions are grouped as “strong” (with absolute strength 1) and “weak” (with absolute strength *w*), as shown in Table 3. Based on experimental data, feedback from CSD to uPN is strong after food deprivation, hence we modeled this non-synaptic interaction by setting a weight from CSD to uPN to be 2, i.e., *W* (5, 3) = 2 under food deprivation. CSD inhibits pLN0 through serotonin, modeled by parameter *β*. Cobra mPN shows stronger activation in the fed state and thus receives basal input *α*. As seen in Table 3, a neuron receives several inputs: from ORN, pLN, CSD or baseline input. We can write it in the vector notation:

**Table 3.**
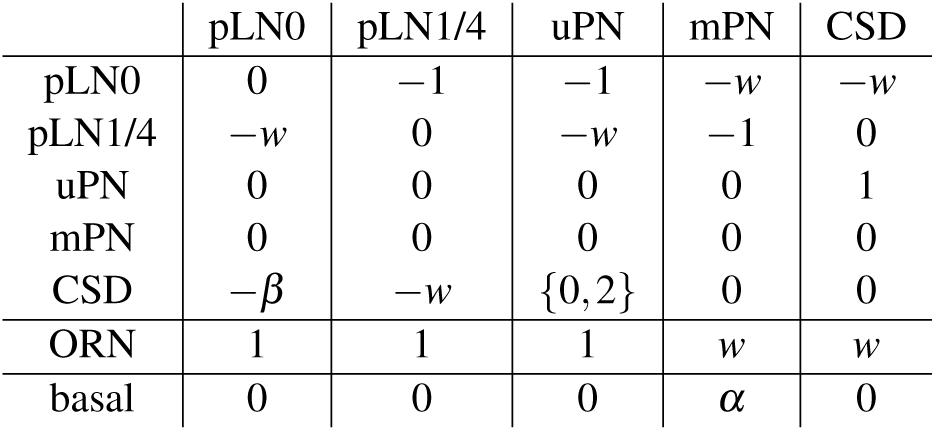
Connection weights (***W***) modeling synaptic and non-synaptic interactions (top five rows), and ORN and basal input of each neuron (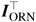 and 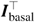 respectively, bottom two rows)

|  | pLN0 | pLN1/4 | uPN | mPN | CSD |
| --- | --- | --- | --- | --- | --- |
| pLN0 | 0 | -1 | -1 | -w | -w |
| pLN1/4 | -w | 0 | -w | -1 | 0 |
| uPN | 0 | 0 | 0 | 0 | 1 |
| mPN | 0 | 0 | 0 | 0 | 0 |
| CSD | $-\beta$ | -w | $\{0, 2\}$ | 0 | 0 |
| ORN | 1 | 1 | 1 | w | w |
| basal | 0 | 0 | 0 | $\alpha$ | 0 |

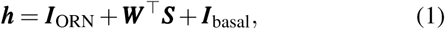

where ***I***_ORN_ is the input vector from ORNs, ***I***_basal_ is the basal input vector, and ***W*** is the interaction matrix between pLNs, PNs and CSD. The *i*th neuron’s state in the next time step is determined by the total input *h*_*i*_ it receives via the following rule:

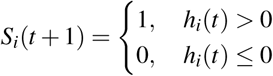

for *i* = 1, 2, 4 (corresponding to pLN0, pLN1/4 and mPN), or

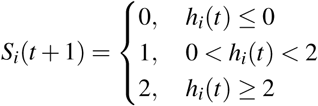

for *i* = 3, 5 (corresponding to uPN and CSD).

To fit our model parameters, we used the steady state activity at the fed state and food-deprived state of the wild-type larvae from experiments, shown in Table 4.

**Table 4.** Activity patterns of neurons in fed and food-deprived state of larvae.

|  | pLN0 | pLN1/4 | uPN | mPN | CSD |
| --- | --- | --- | --- | --- | --- |
| Fed | 1 | 0 | 1 | 1 | 1 |
| Food-deprived | 0 | 1 | 2 | 0 | 2 |

These neural activity patterns are stable and hence impose a constraint on the model parameters *w, α, β* by

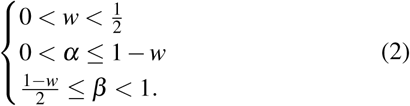

Consideration of perturbation experiments further constrain the upper bound on *β, β <* 1 − *w*. In our model, we choose a set of parameters that fulfills all the constraints: *w* = 0.25, *α* = 0.5, *β* = 0.5.

To relate neural activity to behavioral readout (preference index), we assume a binary readout is determined by

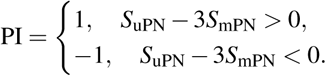

The exact value of the prefactor before *S*_mPN_ does not matter, as long as it is larger than 2. Thus, when mPN is on it generates avoidance behavior, otherwise the fly larvae is attracted by the odor. Our model correctly predicts all the behavioral output of perturbation experiments (Fig. 4C).

## Supplemental figures

**Figure S1.**
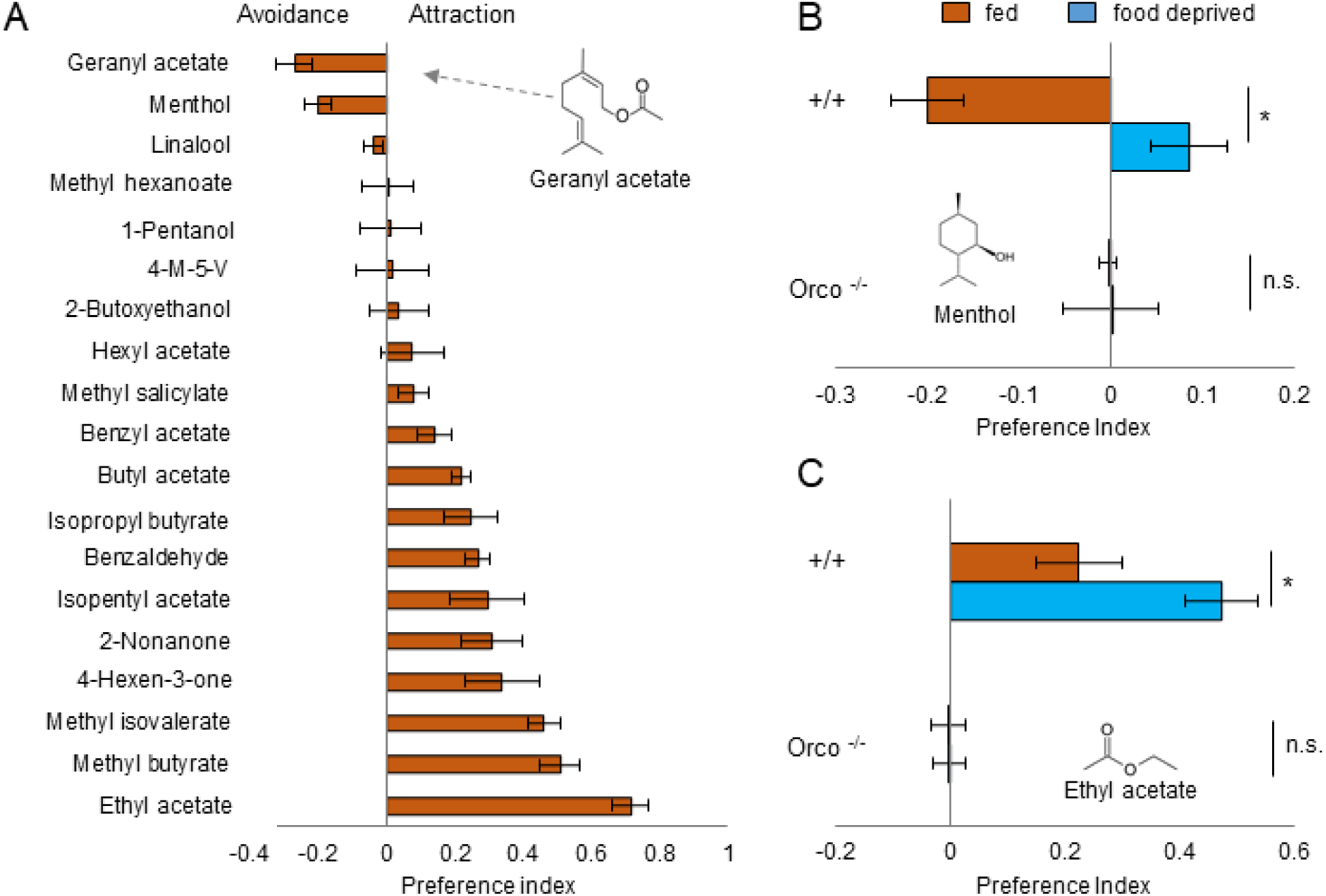
Food deprivation induces a change in olfactory decision making across odorants and is ORN dependent. **A** Odorant screening in fed larvae. Only menthol and geranyl acetate induced aversion, strongest attraction was elicited by ethyl acetate. All odors were tested with 10^−4^ dilution, except menthol (10^−3^). **B** Larvae change their response to menthol (conc. 10^−3^) after food deprivation from avoidance to attraction (two-sample t-test, *p <* 0.001). Mutant larvae (*Orco*^−*/*−^) that lack functional ORNs do not show any significant response to the odor in the fed and food deprived state (one-sample t-test, *p >* 0.05)(*n* = 8). **C** Larvae show increased attraction to ethyl acetate (conc. 10^−6^) after food deprivation (two-sample t-test, *p <* 0.05). Mutant larvae (*Orco*^−*/*−^) that lack functional ORNs do not show any significant response to the odor in the fed and food deprived state (one-sample t-test, *p >* 0.05)(*n* =6–8).

**Figure S2.**
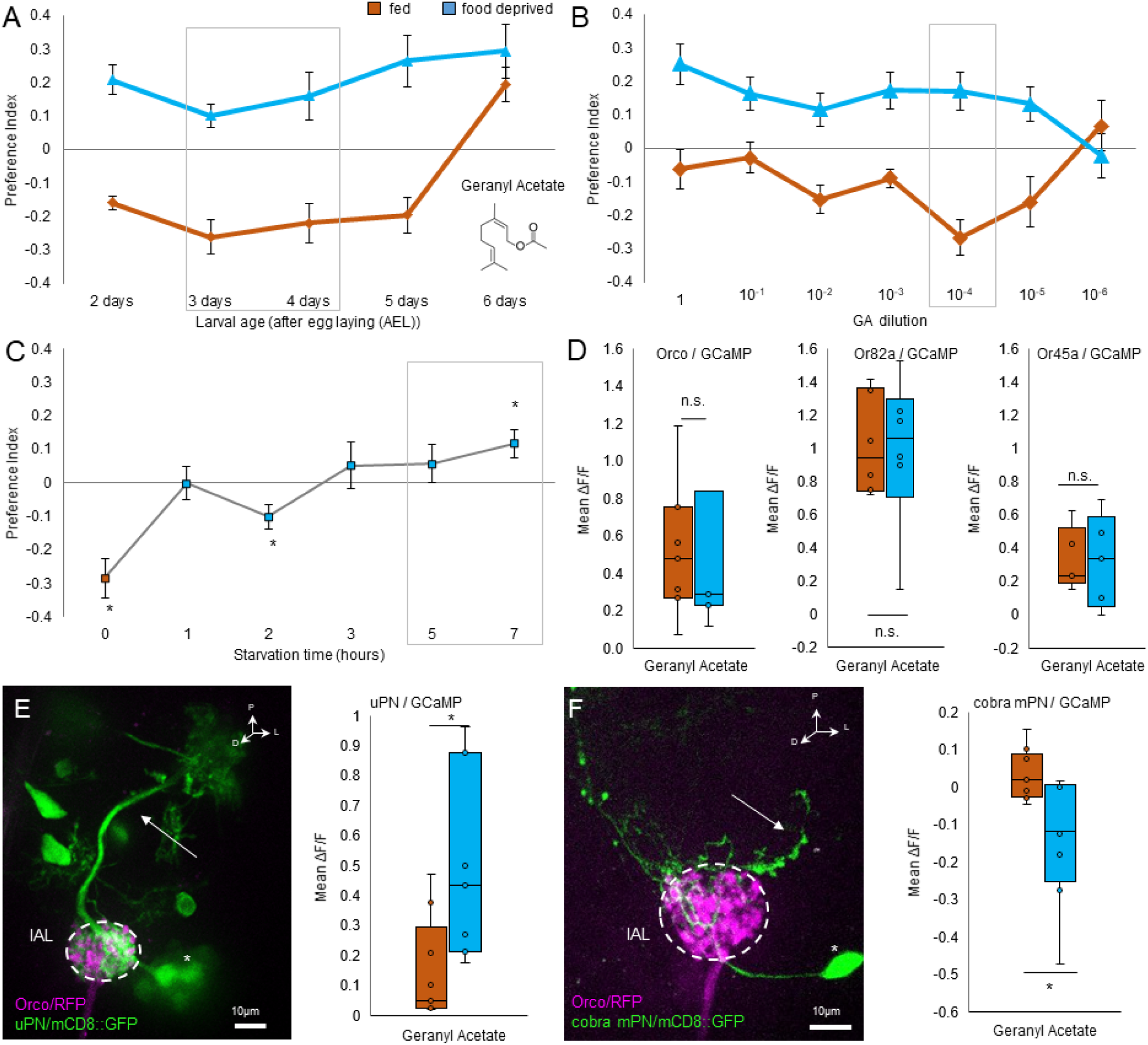
GA behavior is stable across test parameters. **A** The switch in GA 10^−4^ response is present over all feeding stages of the larvae (2-5 days AEL, two-sample t-test, *p <* 0.001). Larvae entering the wandering state show attraction also in the fed state (6 days AEL, two-sample t-test, *p >* 0.05). (*n* = 8–10). **B** For each dilution of GA tested, except the lowest (two-sample t-test, *p >* 0.05), we find a significant switch in behavior between states (two-sample t-test, *p <* 0.05, strongest effect for 10^−4^ : two-sample t-test, *p <* 0.001) (*n* = 6–14). **C** Response to GA 10^−4^ switches to attraction after food deprivation (one-way ANOVA, *p <* 0.001). In fed state larvae avoid the odor (one sample t-test, *p <* 0.001). After short food deprivation larvae lose the avoidance towards GA (1h, 3h, 5h, one sample t-test, *p >* 0.05) or only slightly avoid it (2h, one sample t-test, *p <* 0.05). They are significantly attracted to the odor after 7 hours of food deprivation (one sample t-test, *p <* 0.05). (*n* = 12-16). **D** Calcium activity in the ORNs in response to GA. Mean change in response normalized to baseline before odor presentation. ORNs respond to GA (10^−8^) in fed and food deprived state, however there is no change in response upon food deprivation (two-sample t-test, *p >* 0.05). (*n* = 7). The OR82a neuron shows same GA (10^−6^) response in both states (two-sample t-test, *p >* 0.05) (n=6). The OR45a neuron shows same GA response (10^−6^) in both states (two-sample t-test, *p >* 0.05) (*n* = 5). **E** *GH146-GAL4* labels uniglomerular projection neurons (uPN line). Arrow indicates axonal projection. Asterisk indicates cell bodies. lAL = larval Antennal Lobe. Calcium activity in the uPNs in response to GA. Mean change in response normalized to baseline before odor presentation. The uPNs respond stronger to odors in the food deprived state (two-sample t-test, *p <* 0.05). (*n* = 7–9). **F** *GMR32E03-GAL4* labels the cobra mPN. Arrow indicates axonal projection. Asterisk indicates cell body. lAL = larval Antennal Lobe. Calcium activity in the cobra mPN in response to GA. Mean change in response normalized to baseline before odor presentation. The cobra mPN is inhibited by odors in the food deprived state (two-sample t-test, *p <* 0.05). (*n* = 8–9). Grey boxes indicate parameters used throughout the study. Data points in **A-C** represent pooled data from 5-15 min during testing (mean ± SEM).

**Figure S3.**
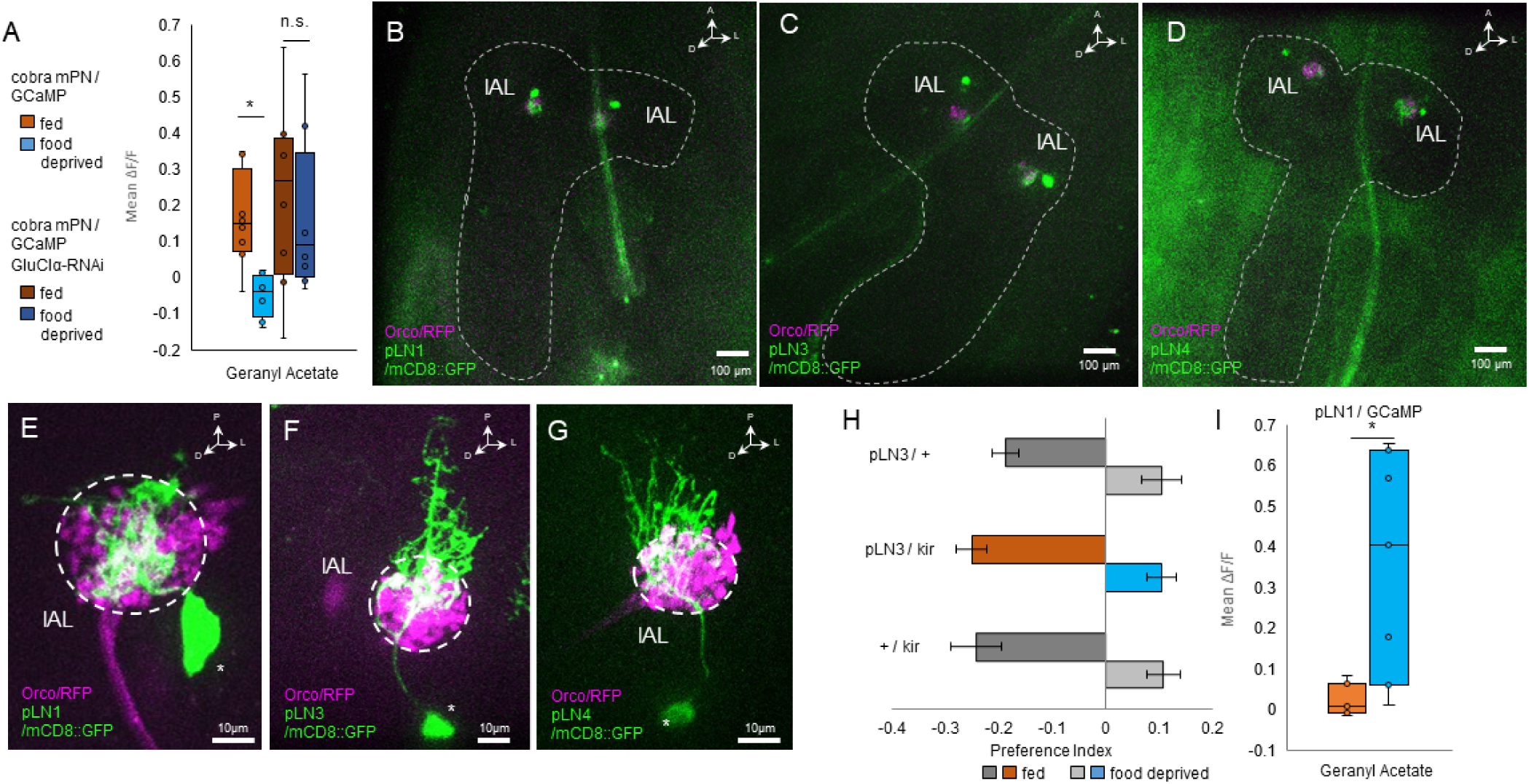
Cobra mPN is inhibited by glutamate released from pLN1,4, but not pLN3. **A** Calcium activity in the cobra mPN in response to GA. Mean change in response normalized to baseline before odor presentation. Light colors: The cobra mPN is inhibited by GA in the food deprived state (two-sample t-test, *p <* 0.01). Dark colors: Upon GluCl*α*-receptor knockdown the cobra mPN shows same response in both states (two-sample t-test, *p >* 0.05).(*n* = 8). **B-D** Whole brain expression patterns of pLN 1, pLN3 and pLN 4 Split-GAL4 lines. Dashed line = larval brain outline. lAL = larval Antennal Lobe. **E-G** Expression patterns of pLN 1, pLN3 and pLN 4 Split-GAL4 lines in the lAL. Asterisks indicate cell bodies. lAL = larval Antennal Lobe. **H** Block of pLN3 neuronal output has no effect on odor responses in both states (one-way ANOVA, *p >* 0.05). (*n* = 8–10) **I** Calcium activity in the pLN1 in response to GA. Mean change in response normalized to baseline before odor presentation. The pLN1 does not respond to GA (10^−6^) in the fed state, but shows increased response after food deprivation (two-sample t-test, *p <* 0.05). (*n* = 7). Bar graphs represent pooled data from 5-15 min during testing (mean ± SEM).

**Figure S4.**
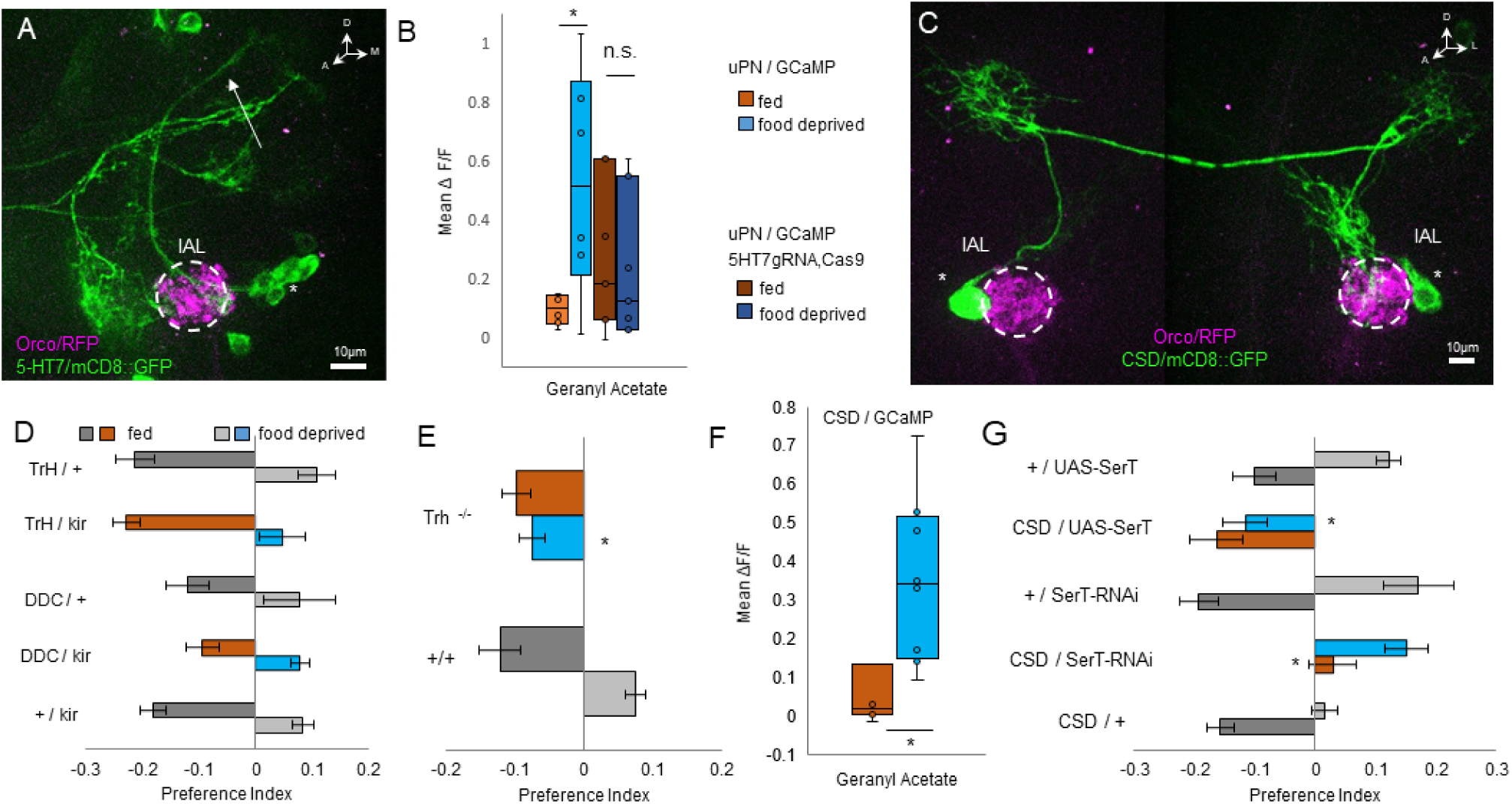
Serotonin released from CSD excites uPNs. **A** Expression pattern of 5-HT7-GAL4. uPNs are labeled in the lAL. Arrow indicates axonal projections. Asterisk indicates cell bodies. lAL = larval Antennal Lobe. **B** Calcium activity in the uPNs in response to GA 10^−6^. Mean change in response normalized to baseline before odor presentation. Light colors: The uPNs show increased response to the odor in the food deprived state (two-sample t-test, *p <* 0.05). Dark colors: Upon knockout of 5-HT7 receptor in the uPNs, they show the same response in both states (two-sample t-test, *p >* 0.05). (*n* = 6–7). **C** *GMR60F02-GAL4* labels the CSD neuron. Asterisk indicates cell bodies. lAL = larval Antennal Lobe. **D** Blocking output of neurons labeled by *Trh-GAL4* does not affect fed (one-way ANOVA, *p >* 0.05) nor food-deprived behavior (one-way ANOVA, *p >* 0.05) towards GA. (*n* = 6–10). Blocking output of neurons labeled by *DDC-GAL4* does not affect fed (one-way ANOVA, *p >* 0.05) nor food-deprived behavior (one-way ANOVA, *p >* 0.05) towards GA. (*n* = 6–14) **E** Mutant larvae (*Trh*^−*/*−^) unable to synthesize serotonin show normal GA avoidance in fed state similar to control levels (two-sample t-test, *p >* 0.05), however they do not switch to GA attraction after food deprivation (two-sample t-test, *p <* 0.01) (*n* = 6–8). **F** Calcium activity in the CSD neuron in response to GA (10^−8^). Mean change in response normalized to baseline before odor presentation. The CSD neuron responds stronger to GA in the food deprived state (two-sample t-test, *p <* 0.05). (*n* = 8–10). **G** Upon knockdown of the serotonin transporter in the CSD neuron using SerT-RNAi we see a significant decrease in odor avoidance in fed larvae (one-way ANOVA, post hoc pairwise comparison, *p <* 0.01) but no effect on food deprived attraction (one-way ANOVA, post hoc pairwise comparison, *p >* 0.05) (*n* = 8–10). Overexpression of the serotonin transporter in the CSD neuron using UAS-SerT leads to a loss of attraction in the food deprived state (one-way ANOVA, post hoc pairwise comparison, *p <* 0.01), however odor avoidance in the fed state is unaffected (Kruskal Wallis test, *p >* 0.05). (*n* = 8). Bar graphs represent pooled data from 5-15 min during testing (mean ± SEM).

**Figure S5.**
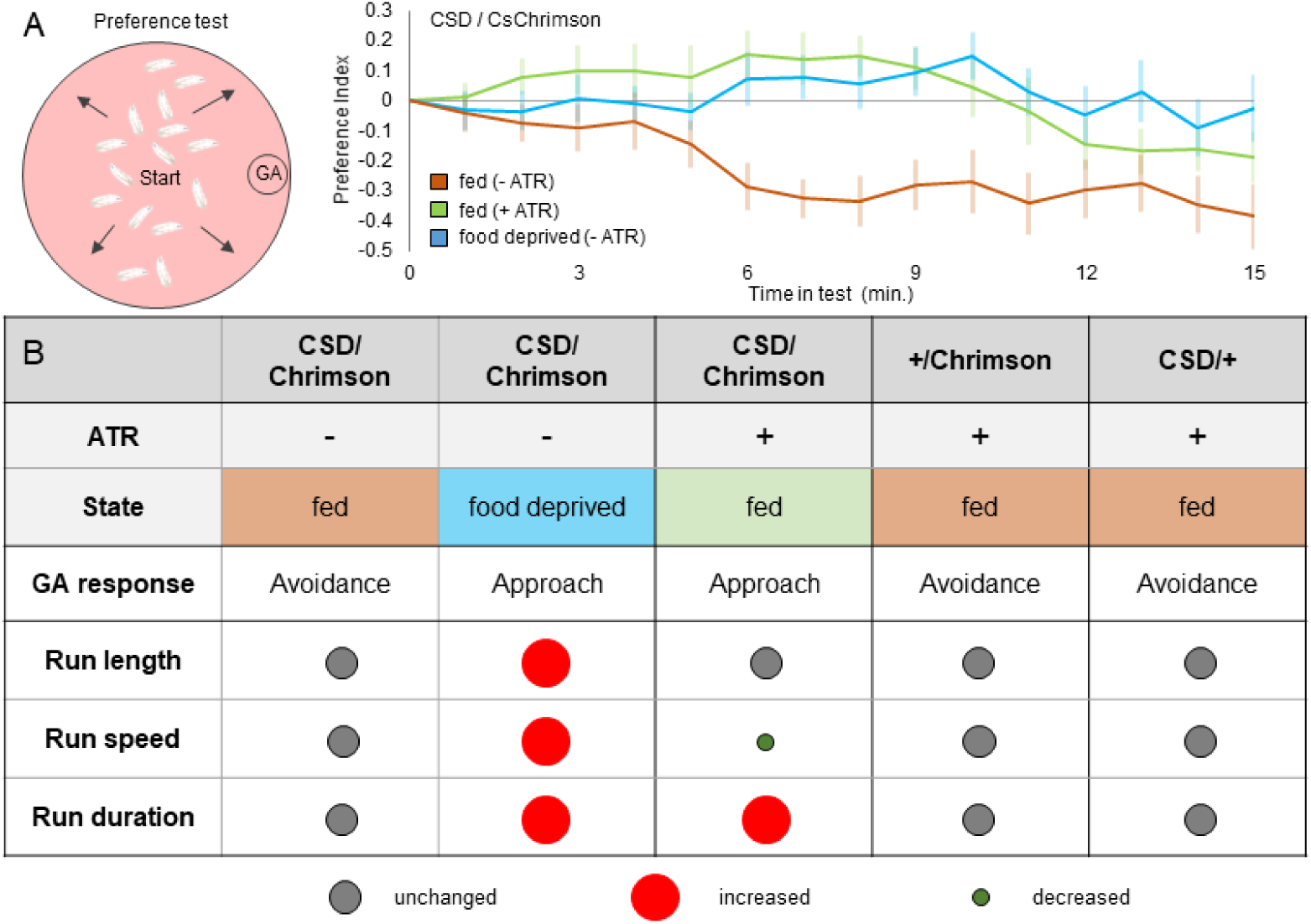
CSD activation affects locomotion parameters differently than food deprivation. **A** Activation of the CSD neuron by expressing UAS-Chrimson in fed larvae mimics the food deprived olfactory response behavior. **B** Locomotion parameters of larvae. Food deprived larvae show increased run length due to increased run speed and run duration. Artificial activation of CSD only induces an increase in run duration.

**Figure S6.**
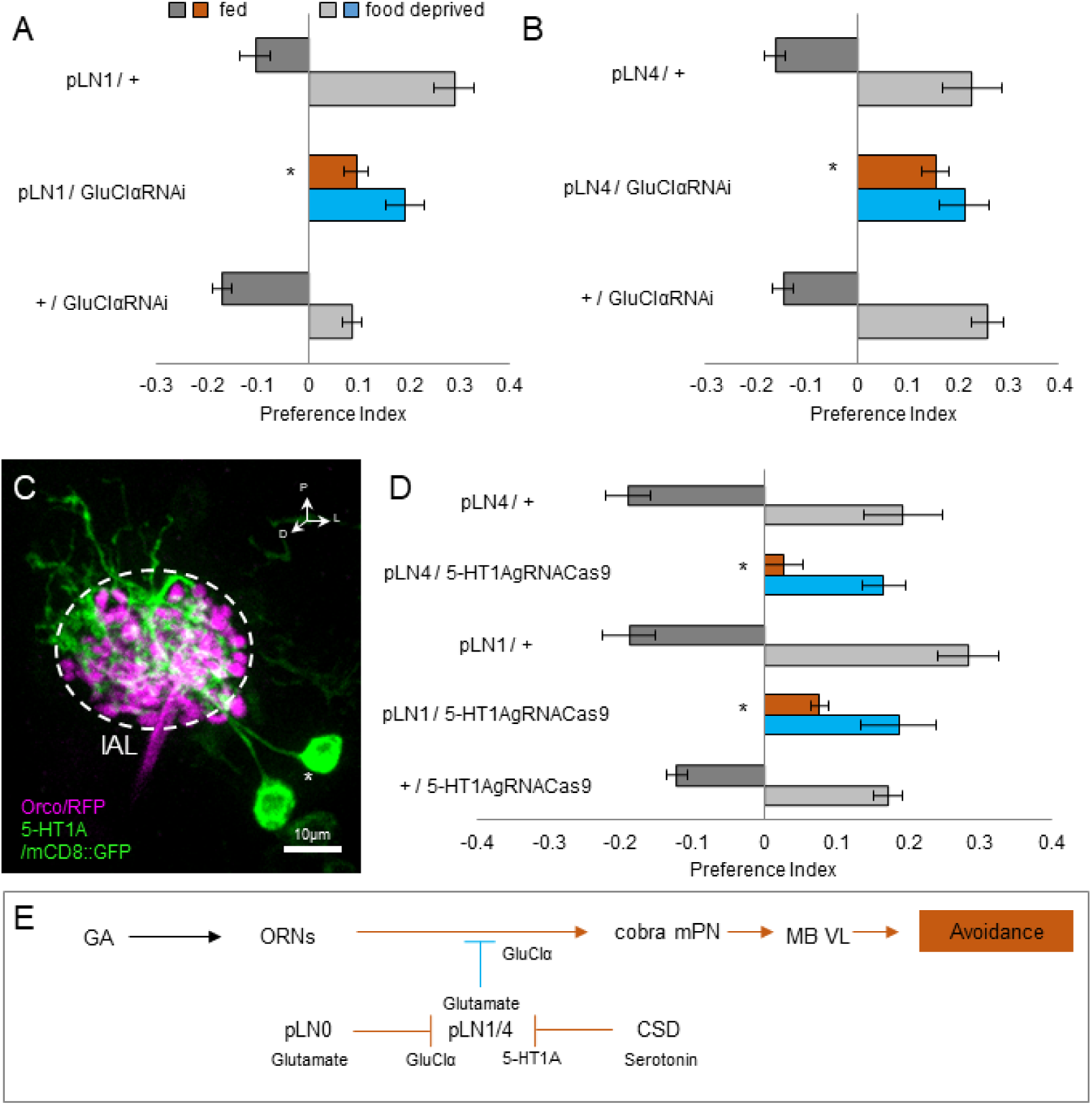
pLN1/4 receive glutamatergic and serotonergic inhibition in the fed state. **A** Knockdown of the GluCl*α*-receptor in pLN1 impairs odor avoidance in the fed state (one-way ANOVA, posthoc pairwise comparison, *p <* 0.001). Odor attraction in the food deprived state is not affected (one-way ANOVA, posthoc pairwise comparison, *p >* 0.05) (*n* = 12–8). **B** Knockdown of the GluCl*α*-receptor in pLN4 neurons impairs odor avoidance in the fed state (one-way ANOVA, posthoc pairwise comparison, *p <* 0.001). Odor attraction in the food deprived state is not affected (one-way ANOVA, *p >* 0.05) (*n* = 7–8). **C** Expression pattern of *5-HT1A-GAL4*. Two local lAL neurons are labeled. Asterisk indicates cell bodies. lAL = larval Antennal Lobe. **D** Knockdown of the 5HT1A receptor in the picky LNs leads to loss of avoidance in the fed state (one-way ANOVA, posthoc pairwise comparison, *p <* 0.001), however attraction in food deprived state is unaffected (one-way ANOVA, *p >* 0.05). (*n* = 6–16). **E** In the fed state, pLN1/4 receive inhibition from a glutamatergic neuron (presumably pLN0) and the serotonergic CSD neuron. Therefore, they are not able to inhibit the downstream cobra mPN. Bar graphs represent pooled data from 5-15 min during testing (mean ± SEM).

**Figure S7.**
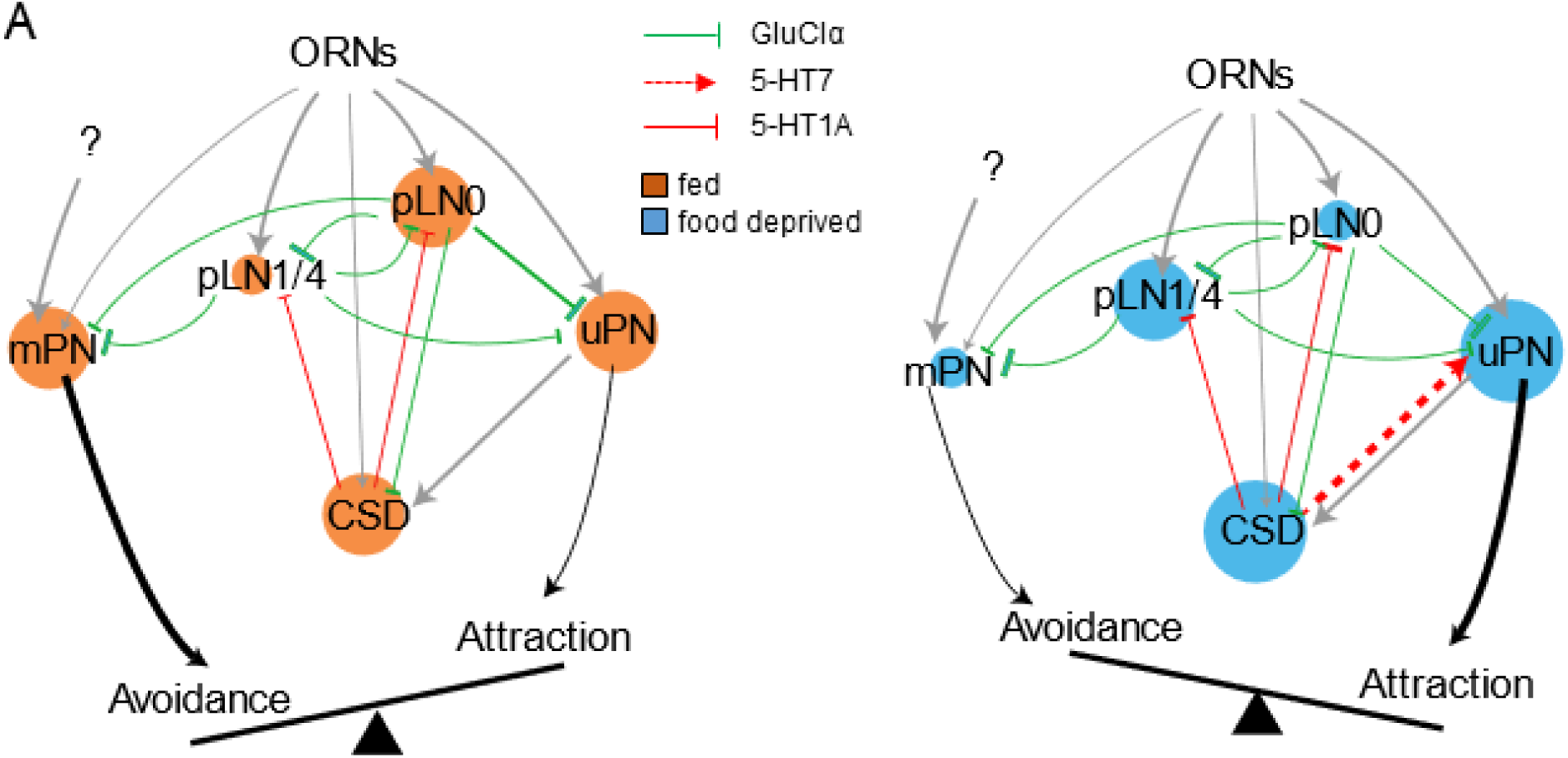
Dynamical systems model connectivity and cell activity. **A** Schematic of model connectivity based on EM data and experimental results. Neuronal activation in the fed (orange) and food deprived (blue) state is indicated in the size of circles.

**Figure S8.**
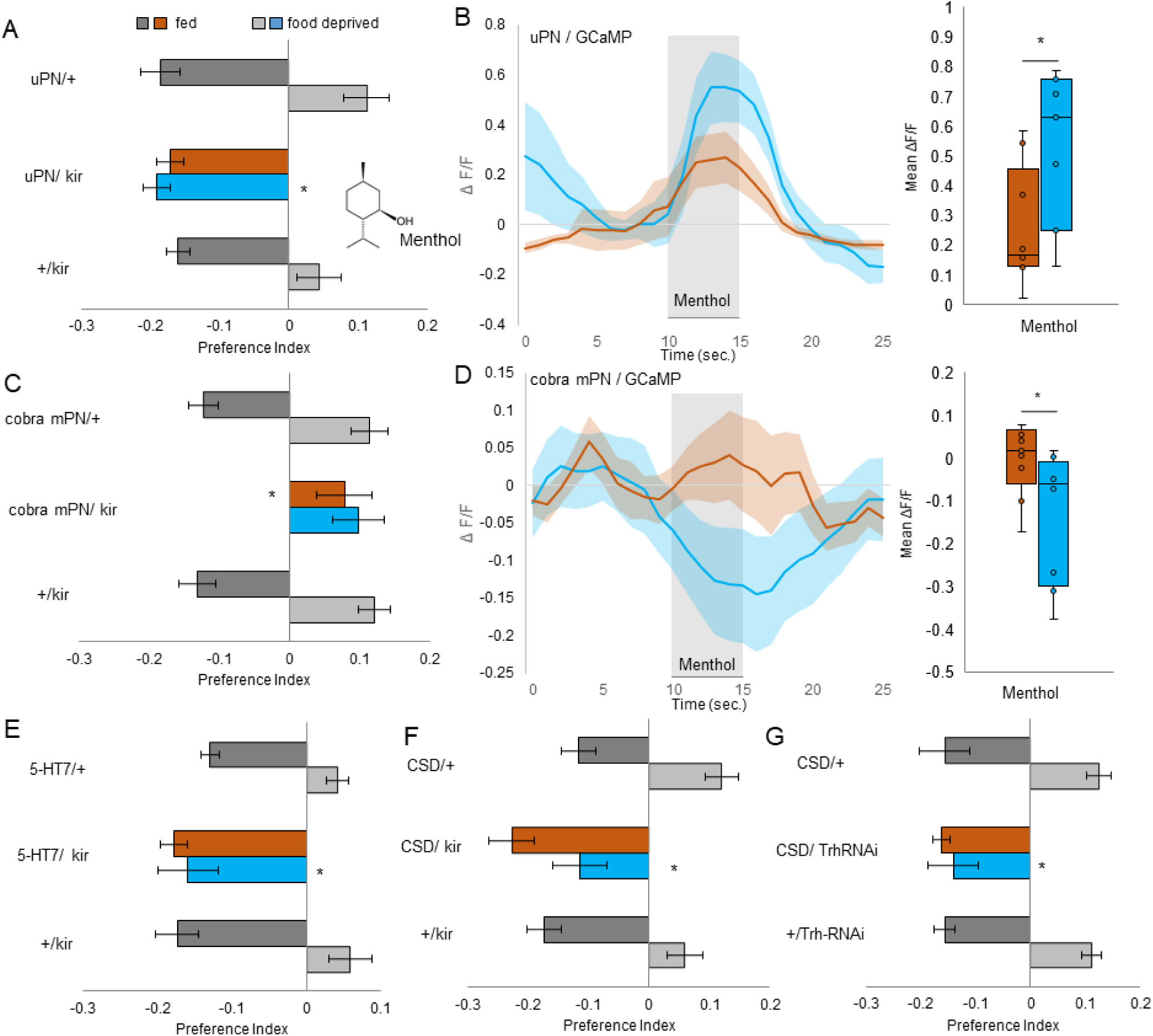
State dependent olfactory behavior in response to the odor menthol requires the same neural circuit as for GA. **A** Blocking output of uPNs labeled by *GH146-GAL4* with *UAS-kir2*.*1* leads to impairment of the food deprived menthol response (one-way ANOVA, posthoc pairwise comparison, *p <* 0.001), but not the fed menthol response (one-way ANOVA, *p >* 0.05). (*n* = 4–6). **B** uPNs show increased response to menthol after starvation (two-sample t-test, *p <* 0.05) (*n* = 7–9). **C** Blocking output of cobra mPN labeled by *GMR32E03-GAL4* leads to impaired odor avoidance in the fed state (one-way ANOVA, posthoc pairwise comparison, *p <* 0.01). However, no effect on odor attraction in the food deprived state can be found (one-way ANOVA, *p >* 0.05). (*n* = 6). **D** After food deprivation, cobra mPN shows decreased response to menthol (two-sample t-test, *p <* 0.05)(*n* = 8–9). **E** Blocking output of neurons labeled by *5-HT7-GAL4* with *UAS-kir2*.*1* leads to impairment of the food deprived menthol response (one-way ANOVA, posthoc pairwise comparison, *p <* 0.01), but not the fed menthol response (one-way ANOVA, *p >* 0.05). (*n* = 8–10). **F** Blocking the output of the CSD neuron with *UAS-kir2*.*1* has no effect in fed state (one-way ANOVA, *p >* 0.05), but leads to impaired food deprived behavior towards menthol (one-way ANOVA, posthoc pairwise comparison, *p <* 0.01). (*n* = 8–10) **G** Knockdown of serotonin synthesis in the CSD neuron does not affect menthol avoidance in fed state (one-way ANOVA, *p >* 0.05). In food deprived state we see an effect compared to controls (one-way ANOVA, posthoc pairwise comparison, *p <* 0.01). (*n* = 8). Bar graphs represent pooled data from 5-15 min during testing (mean ± SEM).

